# The primary cilium dampens proliferative signaling and represses a G2/M transcriptional network in quiescent myoblasts

**DOI:** 10.1101/833061

**Authors:** Nisha Venugopal, Ananga Ghosh, Hardik Gala, Ajoy Aloysius, Neha Vyas, Jyotsna Dhawan

**Affiliations:** CSIR-Centre for Cellular and Molecular Biology, Hyderabad, 500007 India; Institute for Stem Cell Science and Regenerative Medicine, Bengaluru, 560065 India; St. John’s Research Institute, Bengaluru, 560034 India; Academy of Scientific and Innovative Research, Ghaziabad, 201002, India; National Center for Biological Sciences, Bengaluru, 560065, India

**Keywords:** Quiescence, myoblasts, G0, primary cilium, signaling

## Abstract

Reversible cell cycle arrest (quiescence/G0) is characteristic of adult stem cells and is actively controlled at multiple levels. G0 cells extend a primary cilium, which functions as a signaling hub, but how it controls the quiescence program is not clear. Here, we report that primary cilia distinguish different states of cell cycle exit: quiescent myoblasts elaborate a primary cilium *in vivo* and *in vitro*, but terminally differentiated myofibers do not. Myoblasts where ciliogenesis is ablated using RNAi against a key ciliary assembly protein (IFT88) can exit the cell cycle but display an altered quiescence program and impaired self-renewal. Specifically, the G0 transcriptome in IFT88 knockdown cells is aberrantly enriched for G2/M regulators, suggesting a focused repression of this network by the cilium. Cilium-ablated cells also exhibit features of activation including enhanced activity of Wnt and mitogen signaling, and elevated protein synthesis via inactivation of the translational repressor 4EBP1. Taken together, our results show that the primary cilium integrates and dampens proliferative signaling, represses translation and G2/M genes, and is integral to the establishment of the quiescence program.

**Summary statement:** The primary cilium contributes to reversible arrest (quiescence) in skeletal muscle myoblasts, by coordinating and dampening mitogenic signaling focused on a G2/M transcriptional program and protein synthesis.

## Introduction

Primary cilia are microtubule-based membrane-encased organelles that function to receive and transduce signals from the extracellular milieu (Singla and Reiter, 2006). These cellular antennae are intimately linked to the cell cycle, and barring rare instances (Paridaen et al., 2013; Riparbelli et al., 2012), the extension of the cilium is restricted to cells in G0/G1, and suppressed in other cell cycle phases (Goto et al., 2017). The primary cilium is formed from and anchored by a basal body, which is essentially a sequestered centrosome that has been modified for this role. Upon cell cycle re-entry, cilia are actively dismantled prior to mitosis, releasing the basal body to function as a microtubule-organizing centre (MTOC) for spindle assembly. Multiple proteins that play important ciliary roles are also involved in cell cycle regulation: for example, Aurora Kinase A (AurA), Polo Like Kinase-1 (Plk1), and Anaphase Promoting Complex (APC) are all key regulators of centrosome function and ciliary disassembly, as well as mitotic progression (reviewed in (Walz, 2017)). Thus, the current understanding is that the cilium exerts a check on cell cycle progression (reviewed in (Goto et al., 2013)) and that disassembly of the cilium is a prerequisite for cell cycle re-entry (Kim et al., 2011). The cilium is vital in embryonic patterning (Hirokawa et al., 2006), and its role in terminally differentiated adult cells such as kidney epithelia is critical for their function (Yoder, 2007). Mutations in ciliary genes give rise to a spectrum of complex diseases collectively termed ciliopathies (Hildebrandt et al., 2011), usually associated with cellular overgrowth, thus supporting an inhibitory influence of cilia on proliferation. However, the molecular mechanisms that couple ciliogenesis and cell cycle progression are still emerging and the functions of the cilium within the quiescent cell are largely unexplored.

The quiescent state is critical for the function of adult stem cells, which contribute to regeneration and tissue homeostasis (reviewed in (Rumman et al., 2015)). In skeletal muscle, homeostatic maintenance of differentiated tissue, as well as its regeneration following injury, are made possible by a small population of stem cells called muscle satellite cells (MuSC) (Mauro, 1961). These muscle precursors persist in a quiescent undifferentiated state in adult muscle and are activated to re-enter the cell cycle when the tissue experiences damage. The majority of activated MuSC enter a program of proliferation and differentiation to regenerate damaged myofibers, while a minor subset enters a distinct program to self-renew and regenerate myofiber-associated quiescent MuSC, thus leading to functional tissue recovery (Morgan and Partridge, 2003). Although a number of signaling, metabolic, transcriptional and epigenetic regulatory circuits have been implicated in MuSC quiescence (Cheung and Rando, 2013), the mechanisms that govern its establishment during early postnatal development and its maintenance during adult life are still being uncovered. Recently, quiescent MuSC were shown to possess primary cilia, which were implicated in the regulation of asymmetric cell division during cell cycle activation (Jaafar Marican et al., 2016). However, the genetic and signaling networks potentially controlled by the cilium within quiescent MuSC are not known.

Here, we investigate the role of the primary cilium in quiescence using an established culture system to model alternate MuSC fates: asynchronously proliferating C2C12 myoblasts can be driven to enter either reversible (Arora et al., 2017; Sachidanandan et al., 2002) or terminal cell cycle exit (Blau et al., 1983), by modulating culture conditions. We show that not only are primary cilia present on quiescent MuSC *in vivo*, and elaborated during quiescence *in vitro*, but are also transiently induced during early myogenic differentiation. When ciliogenesis is ablated using RNAi against IFT88, a key intraflagellar transport protein involved in ciliary extension, an altered quiescence-reactivation program is observed. Additionally, IFT88 knockdown in G0 leads to specific induction of G2/M transcriptional networks, and cells lacking cilia display enhanced activity of multiple cilium-associated signaling pathways, accompanied by an increase in protein synthesis, with a specific shift towards 4EBP1-regulated translation. Intriguingly, knockdown cells do not show increased DNA synthesis or mitosis. These observations suggest that in quiescent cells, loss of this cellular antenna relieves the repression on mitogenic signaling but additional events are required to trigger the exit from quiescence. Our findings support the existence of complex interplay between the cilium cycle and the cell cycle at both transcriptional and translational levels. Overall, our study reveals that specific repression of G2/M genes by the primary cilium is integral to the establishment of the self-renewing quiescent state in culture, with implications for stem cell maintenance and function.

## Results

### Primary cilia are dynamically regulated during the cell cycle in myogenic cells

Earlier, we showed that quiescent myoblasts in culture display a distinct transcriptional profile, with altered signaling modules, in particular, enhanced Wnt-TCF signaling (Aloysius et al., 2018; Subramaniam et al., 2013). Interestingly, a number of cilium-associated genes showed increased expression in G0. We compiled a putative “ciliome” consisting of 1896 cilium-related genes curated from available datasets from different cell types (Kim et al., 2010; McClintock et al., 2008). Comparison of the “ciliome” with the “G0 altered” transcriptome consisting of 1747 annotated Differentially Expressed Genes derived from a microarray analysis of C2C12 myoblasts in proliferation vs. quiescence (Subramaniam et al., 2013), yielded substantial overlap (345 genes [19%], *p*-value: 1.717854e-19) (Fig. S1A). Selected transcripts encoding cilium-associated proteins were validated as up-regulated in quiescent myoblasts (Fig. S1B). Together, these findings suggest a previously unreported induction of a ciliary transcriptional program in skeletal muscle myoblast quiescence.

To investigate whether the transcriptional induction of ciliary genes is accompanied by the elaboration of primary cilia in G0 myoblasts, we used immunofluorescence analysis. C2C12 myoblasts grow asynchronously in adherent culture and can be triggered to enter reversible arrest (quiescence/G0) by culture in suspension, where deprivation of adhesion leads to cessation of cell division (Milasincic et al., 1996; Sachidanandan et al., 2002). G0 is rapidly reversed upon re-plating onto a culture surface, wherein cells synchronously re-enter the cell cycle. In actively proliferating cultures, 24% of cells were found to be ciliated (Fig. 1A, B). This population heterogeneity is consistent with the asynchronous distribution in different cell cycle phases and the extension of primary cilia only during a restricted window of the cell cycle, in G1/G0. Suspension-arrested myoblasts showed an increase in ciliation with ∼60% of G0 cells marked by primary cilia, which is reversed upon reactivation into the cell cycle, wherein the frequency of ciliated cells returned to 23% within 24 hrs (Fig. 1A, B). In addition, G0 cells showed an increased average ciliary length: 70% of cilia were > 2 μ (Fig. 1C). By contrast, proliferating cells display shorter cilia: only ∼30% of cilia were > 2 μ (Fig. 1C).

**Fig. 1.**
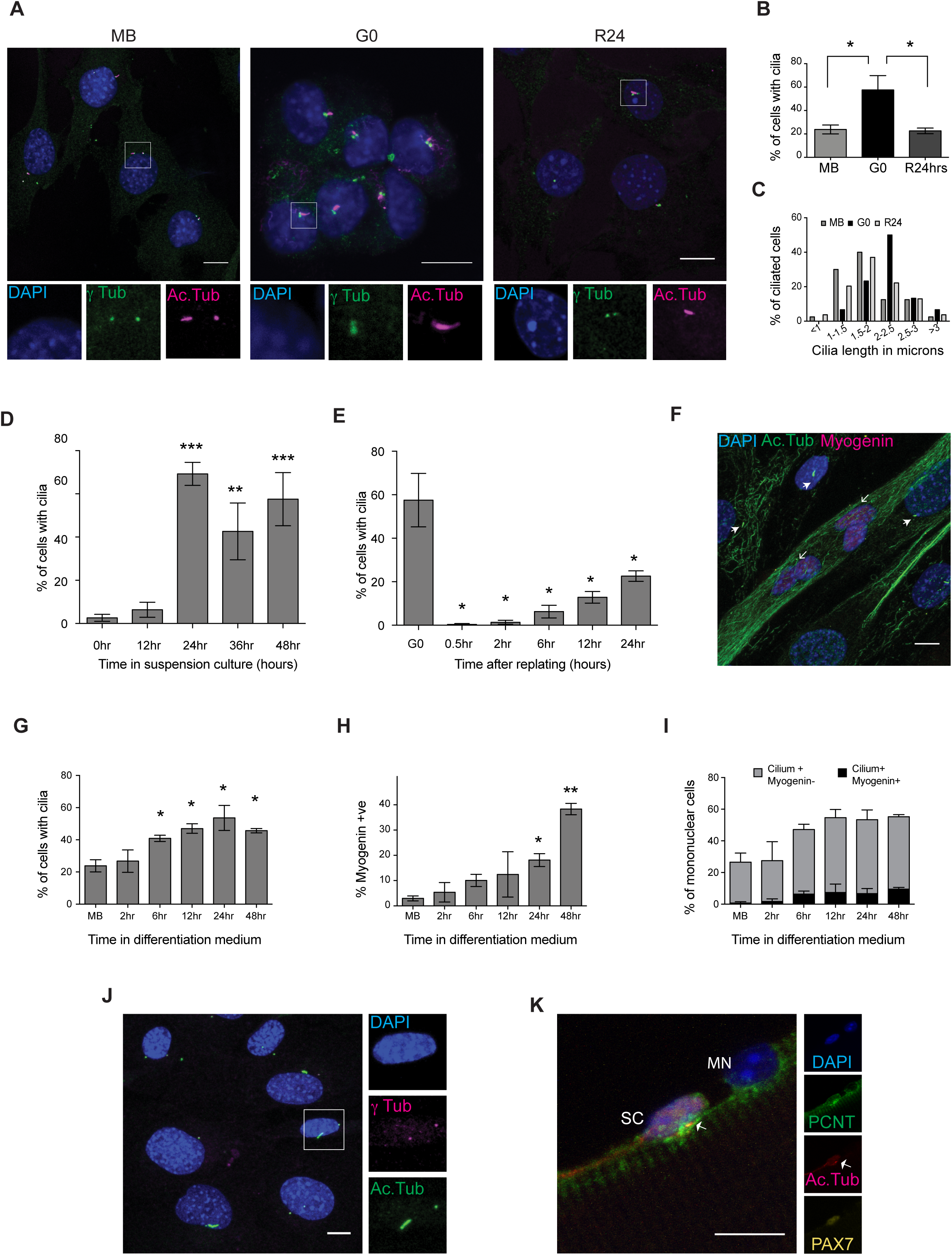
Primary cilia are dynamically regulated and are preferentially present on quiescent myoblasts *in vitro* and *in vivo*. A. Immuno-detection of primary cilia on C2C12 myoblasts in different cellular states - proliferating (MB), quiescent (G0), and 24 hours after reactivation into cell cycle from quiescence (R24). Antibodies against γ-tubulin (γ-tub) mark centrosome and Acetylated tubulin (Ac.Tub) mark cilia; DAPI was used to label DNA. While only a small proportion of MB is ciliated, G0 myoblasts show a higher frequency of ciliation, which is reversed at R24. Lower panel shows magnified views (boxed insets). (Scale bar, 10 μm.) B. Quantification of data showed in (A). Values are mean ± s.e.m, N≥3, * *p* < 0.05 C. The length of cilia present on MB, G0, and R24 cultures was measured and the distribution plotted. Cilia elaborated by G0 myoblasts are longer (Average length 2.2 μm) than those in proliferating cultures (Average length 1.8 μm). The numbers of ciliated cells analyzed are MB (40), G0 (30), and R24 (54). D. Increased frequency of ciliated cells observed as proliferating myoblasts withdraw into quiescence during a time course of suspension arrest (0-48 hours). Values are mean ± s.e.m, N≥3, * *p* < 0.05, ** *p* < 0.01, *** p < 0.005 E. Rapid loss of cilia during reactivation of G0 cells by re-plating on to an adhesive substratum, and gradual increase in ciliation during cell cycle re-entry. Values are mean ± s.e.m, N=3 * *p* < 0.05 F. Immunofluorescence analysis of cilia in 5 day differentiated culture of C2C12 cells: cilia are marked with Acetylated tubulin (Ac.Tub), Myogenin marks differentiated cells. DNA is labeled with DAPI. (Scale bar, 10 μm). Cilia are absent on multinucleated myotubes (open arrowheads), and preferentially present on mononuclear (unfused) cells that lack Myogenin (closed arrowheads). G. Quantitative analysis of ciliation during a time course of differentiation (represented by image in F) reveals increased proportion of ciliated cells as myoblasts fuse to form myotubes. Values are mean ± s.e.m, N=3, * *p* < 0.05 H. Increased proportion of Myogenin**^+^** cells during the period where ciliation was analyzed in (G) highlights the time course of increase in Myogenin expression during differentiation. Values are mean ± s.e.m, N=3, * *p* < 0.05 I. Quantitative analysis of cells that are both ciliated and Myogenin**^+^** shows that ciliated cells are largely negative for Myogenin, indicating exclusion from differentiation. Values are mean ± s.e.m, N=2 J. Primary cilia are present on 47% of reserve cells in differentiated cultures. 5-day differentiated C2C12 cultures were mildly trypsinised to remove myotubes, enriching the adherent undifferentiated mononuclear reserve cells. Cilia were detected using γ-tubulin (γ tub) and Acetylated tubulin (Ac.Tub). (Scale bar, 10 μm) K. Primary cilia are associated with 71.1% Pax7**^+^** satellite cells (SC) but not with Pax7**^-^** myonuclei (MN). Representative immunofluorescence image of a single fiber isolated from mouse EDL muscle and stained for Pax7 to mark satellite cells, and Pericentrin (PCNT) and Acetylated tubulin (Ac.Tub) to mark primary cilia. DAPI marks nuclei. (Scale bar, 10 μm)

To establish the dynamics of cilium acquisition during cell cycle exit, we isolated cells at different time points after induction of quiescence and stained for acetylated-tubulin. Cells showed a rapid increase in ciliation during the early stages of quiescence entry to reach a maximum of ∼60% ciliated cells at 24 hrs post suspension, after which there was little change (Fig. 1D) until the time of harvest (48 hrs). The increase in ciliated cells correlates with the loss of markers of proliferation Ki67 and Cyclin A2, and induction of the growth arrest-specific (GAS) gene PDGFRα (Schneider et al., 2005) (Fig. S2). We also analyzed the dynamics of cilium loss during reactivation out of G0, and observed a rapid and complete loss of primary cilia within 30 minutes of re-plating. With progression past the G0-G1 transition (6 hrs) and entry into S phase (12 hrs), the frequency of ciliated cells gradually increased, reaching ∼20% at 24 hours post re-plating (Fig. 1E), by which time the culture resembles a pre-quiescent population of asynchronously proliferating myoblasts (Fig 1B). These results establish that as in other cell types, quiescence in myoblasts is associated with a reversible extension of longer cilia on a larger proportion of cells than observed in proliferating cultures. The increased frequency and length of cilia in G0 myoblasts, and the rapid loss of ciliation within minutes of reactivation supports the notion of the cilium as an inhibitory influence on cell cycle re-entry and progression, as described in other cell types (Kim et al., 2011).

### Myogenic differentiation involves a transient ciliation event

Myoblasts exit the cell cycle irreversibly during myogenic differentiation. We used immunofluorescence analysis to establish the status of ciliation during entry into terminal arrest. By contrast to reversibly arrested myoblasts, terminally differentiated myotubes lacked primary cilia (Fig. 1F). However, we observed a transient phase of ciliation in early differentiating cultures prior to myoblast fusion, with a steady increase in the frequency of ciliated cells from as early as 6 hrs in differentiation medium, to a maximum of 53% observed at 24 hrs. High levels of ciliation were still observed at 48 hrs when fusion of myoblasts into myotubes was evident (Fig. 1G). A recent report of this transient ciliation suggested that Hh signaling during the induction of myogenic differentiation is mediated by the primary cilium and that deregulation of this node is associated with tumorigenesis (rhabdomyosarcoma formation) (Fu et al., 2014). Interestingly, although the time window of transient ciliation corresponds to the increase in expression of Myogenin, a key regulator of differentiation (Fig. 1H), we found little overlap between cells expressing Myogenin and those bearing a primary cilium in differentiating cultures (Fig. 1I). Our observations suggest that establishment of the myogenic program is specifically excluded in cells bearing primary cilia.

### Reversibly arrested reserve cells in differentiating cultures are ciliated

Cultures triggered for myogenic differentiation yield a heterogeneous population: at 5 days post serum withdrawal, while ∼80% of nuclei may be found in multinucleated myotubes, the remaining ∼20% constitute mononuclear “reserve cells” (Yoshida et al., 1998), a pool of quiescent undifferentiated cells that retain the ability to re-enter the cell cycle if proliferative conditions are restored (Abou-Khalil et al., 2013). We separated out the mononucleated (undifferentiated) fraction from differentiating cultures and verified that these cells do not express Myogenin (Fig. S3). As seen in suspension-arrested G0 myoblasts, ∼50% of unfused reserve cells are ciliated (Fig. 1J) and account for the number of ciliated cells observed in early triggered cultures. Taken together, these results show that the presence of primary cilia distinguishes reversibly arrested from irreversibly arrested cells.

### Quiescent satellite cells *in vivo* are marked by primary cilia

Recently, primary cilia were shown to be preferentially present on quiescent MuSC and not on activated MuSC, where they play a role in asymmetric cell division during the early regenerative response (Jaafar Marican et al., 2016). To investigate whether cilia are associated with reversibly versus irreversibly arrested muscle cells *in vivo*, we isolated single muscle fibers from the mouse Extensor Digitorum Longus (EDL) muscle and probed for the cilium by immunofluorescence analysis. We detected primary cilia on ∼70% of Pax7**^+^** quiescent MuSC but no cilia were found on the rest of the myofiber membrane, either associated with myonuclei (Pax7**^-^**) (Fig. 1K) or elsewhere. Thus, as seen in cultured myoblasts, most quiescent MuSC are ciliated, while the differentiated myofibers are not, indicating that the cilium distinguishes reversible from irreversible arrest *in vivo* as well.

### Abrogation of ciliary extension leads to loss of canonical features of quiescence

Ciliary extension critically depends on intra-flagellar transport (IFT), the process by which material is transported on stable microtubule tracks in the cilium core. IFT is regulated by a number of genes, primarily encoding motor proteins and adaptors that link cargo to the cytoskeleton (Hao and Scholey, 2009). Since ciliary extension was triggered in myoblasts early after receiving cues for cell cycle exit, we postulated that the primary cilium is involved in the regulation of cell cycle exit programs and cell fate determination. To explore this connection, we investigated the ability of cells to enter G0 when ciliogenesis was blocked by siRNA-mediated suppression of a key anterograde transport protein IFT88. siRNA treatment resulted in a 60% reduction in the mRNA levels (Fig. 2A) and a 92% reduction in protein levels of IFT88 (Fig. 2B), compared to control. Knockdown of IFT88 led to significant suppression of ciliogenesis (Fig. 2C, D). Interestingly, we observed a stage-specific effect of cilium knockdown. Contrary to earlier reports where blocking ciliogenesis in HeLa cells results in hyper-proliferation (Robert et al., 2007), cilium knockdown in myoblasts under growth conditions, did not lead to increased proliferation, as evidenced by unchanged frequencies of Ki67^+^ cells (Fig. 2E). However, when knockdown cells were triggered to enter quiescence, we observed an increased frequency of Ki67^+^ cells compared to control cells (Fig. 2F). Knockdown populations also showed a decreased frequency of cells expressing the quiescence marker, p27 (Fig. 2G). Interestingly, cell cycle analysis revealed that myoblasts lacking cilia display an altered FACS profile with an increased proportion of G2/M cells (Fig. 2H). These results indicate that suppressing ciliary extension compromises entry into G0.

**Fig. 2.**
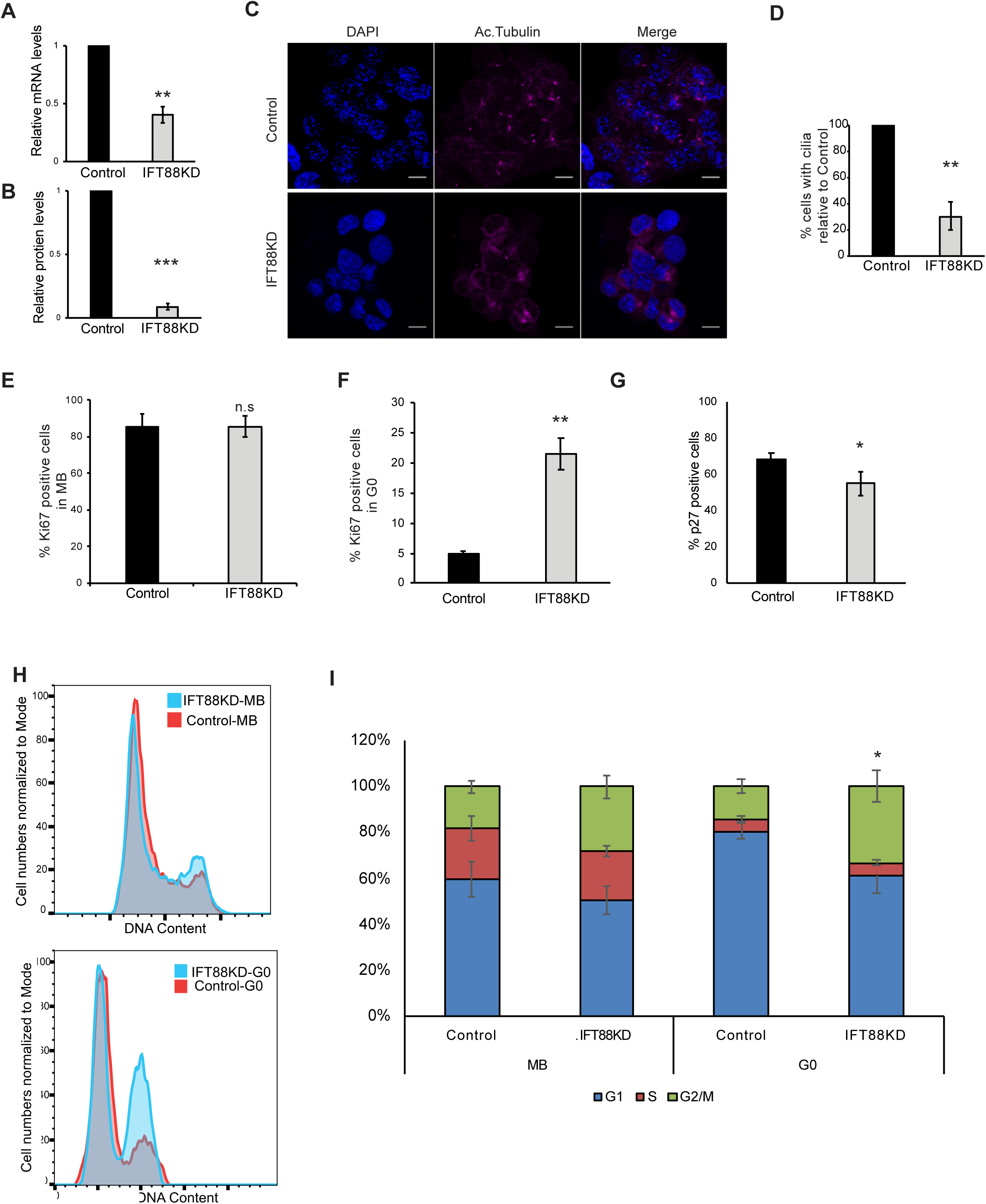
Abrogation of primary cilia has distinct effects in proliferating and quiescent myoblasts. C2C12 myoblasts were transfected with siRNAs targeting IFT88 to block ciliogenesis, and were analyzed for effects of knockdown on proliferation and quiescence. Non-targeting siRNA was used as control. A. qRT-PCR analysis was used to detect levels of IFT88 mRNA in control and siRNA treated samples. Knockdown myoblasts show reduced levels of IFT88 mRNA. Values represent values indicated are mean ± s.e.m, N=3, **p-value = 0.0062. B. Western blotting analysis demonstrates efficient knockdown of respective target protein levels. Values represent values indicated are mean ± s.e.m, N=3, ***p-value= 0.0004. C. Primary cilia were visualized by immunofluorescence labeling of Acetylated tubulin (Ac. tubulin) in IFT88 knockdown myoblasts cultured in quiescence-inducing conditions. Representative immunofluorescence images are shown. A single cilium is present per cell in control cultures, whereas knockdown cells show greatly reduced cilia (accompanied by increased tubulin staining in the cell body). D. Knockdown of IFT88 caused reduction in frequency of ciliated cells. Values represent mean ± s.e.m, N=4, ** p-value = 0.0034. E. Proliferation of cells was analyzed by quantification of Ki67**^+^** cells, detected by immuno-staining. Targeting ciliogenesis had little effect in proliferating conditions (MB). Values represent mean ± s.e.m, N=3, ns = not significant (p-value = 0.959). F. When induced to enter quiescence (G0), knockdown cells displayed increased frequency of Ki67**^+^** compared with control siRNA-transfected cells. Values represent mean ± s.e.m, N=5, ** p-value = 0.0015. G. Ablation of ciliogenesis suppresses expression of quiescence marker p27. C2C12 myoblasts stably expressing the p27 sensor (mVenus-p27K^-^) were transfected with siRNA targeting IFT88or control siRNA and then cultured in quiescence-inducing conditions. The frequency of p27 (mVenus) positive cells was measured by flow cytometry. Values represent mean ± s.e.m, N=3, * p-value = 0.0234. H. Flow cytometric analysis of IFT88 siRNA-transfected cells cultured in conditions that allow proliferation (MB) does not show a significant shift in cell cycle profile. When placed in conditions that induce quiescence (G0) in control cells, knockdown cells display an increased proportion of cells in G2/M. Representative profiles are shown where the X axis shows the DNA content estimated by florescence intensity of the DNA dye DR, and the Y axis represents cell numbers as a percentage of maximum (normalized to mode). I. Quantification of experiment described in (H) showing shifts in proportion of cells in different cell cycle stages (G1, S, G2/M). Values represent mean ± s.e.m, N=3, * p-value= 0.043.

### Transcriptome profiling reveals a stage-specific cell cycle block in IFT88 knockdown cells

To determine the scale of changes that contribute to the altered quiescent state, we used global transcriptome profiling. Proliferating myoblasts (MB) were transfected with either IFT88 siRNA or a non-targeting control siRNA following which cells were transferred to suspension culture for 48 hours to induce quiescence (G0) and subsequently reactivated for 2 hours by re-plating (R2). Affymetrix microarrays were queried with RNA isolated from cells in these three different states and a comparative analysis was performed between control and knockdown samples for each cell state.

The transcriptional profile of IFT88KD myoblasts was consistent with the stage-specific cell cycle phenotype observed and showed maximum shifts from the profile of control cells at conditions of G0 (Fig. 3A, B). Fewer genes were altered in proliferating conditions (51 up-regulated, 31 down-regulated), whereas, knockdown cells held in conditions that normally induce G0 showed a larger number of genes with altered expression (253 genes up-regulated, 157 genes down-regulated). Thus, IFT88KD myoblasts cultured in suspension have a transcriptional profile distinct from control cells (Fig. 3A). A similarly altered profile was also seen in reactivated cells (R2), where 168 genes were up-regulated and 85 genes down-regulated (Fig. 3A, B). Interestingly, in IFT88KD cells, G0 and R2 profiles were very similar, whereas in control cells, substantial alteration of the transcriptome reflecting the reactivation of the cell cycle program. Overall, these observations suggest that suppression of the cilium leads to a block beyond which cells neither entered nor exited quiescence normally.

**Fig. 3.**
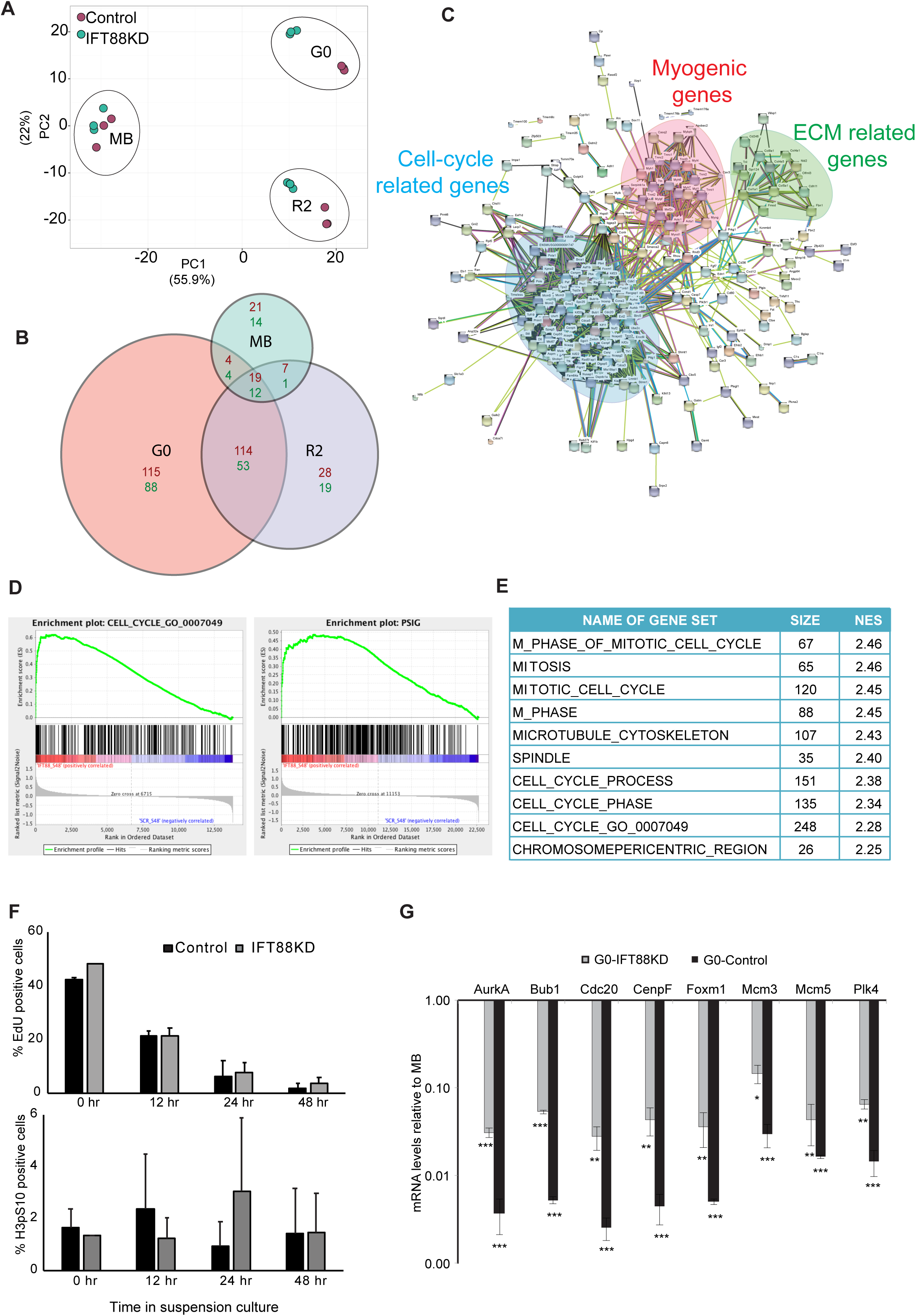
IFT88 knockdown cells display an altered quiescence program enriched with G2/M signature pathways. Affymetrix array-based transcriptional profiling was performed on IFT88 siRNA and control siRNA transfected myoblasts at 3 different cell cycle states -proliferating (MB), quiescent (G0), and 2 hours after reactivation into cell cycle from G0 (R2). A. Two dimensional Principal Component Analysis (PCA) plot (Metsalu and Vilo, 2015) of microarray data showing that IFT88KD G0 cells have a transcriptional profile that is distinct from Control G0 cells. Samples from different cell states can be seen as distinct clusters while the replicates of each sample cluster together indicating that they behave similarly. Control samples are shown as green dots and IFT88KD samples are shown as red dots. Note that control and knockdown cells do not show differences in the MB state, but are well separated in both G0 and R2 states, indicating a stage-specific requirement for the cilium. B. Size adjusted Venn diagram (Hulsen et al., 2008) depicting the number of differentially expressed genes in MB, G0 and R2 states. IFT88KD cells showed the greatest deviation from Control (maximum number of altered genes) in G0. C. STRING analysis was used to visualize the networks formed by the altered genes at G0. Genes up-regulated in IFT88KD cells displayed a strong interaction network, which comprised of three major clusters: (i) Cell cycle-related, (ii) Myogenic genes (iii) ECM-related. Disconnected nodes (genes) are not displayed. D. GSEA was used to identify pathways of genes in the altered transcriptome of IFT88KD G0 myoblasts. The plots show enrichment of genes related to cell cycle and proliferation. The gene sets used as reference here are the 315 genes annotated in the GO term GO:0007049 (CELL CYCLE) and 338 genes identified and collated as a proliferation signature (pSig) E. Gene sets with highest enrichment scores identified in GSEA are specific to G2/M phases. NES represents Normalized Enrichment Score. F. IFT88KD cells show cell cycle exit kinetics similar to Control cells and do not participate in DNA synthesis (EdU incorporation, top panel) or mitosis (H3pS10, bottom panel), during a time course of suspension culture. Values represent mean ± s.e.m, N=2. G. Validation of microarray analysis: qRT PCR analysis was used to determine mRNA levels of selected G2/M genes identified as up-regulated in IFT88KD G0 cells at conditions of proliferation (MB) and quiescence (G0). The relative mRNA levels were calculated in comparison to MB for Control and IFT88KD cells. Control cells show a characteristic repression of expression of mitotic regulators in G0 (varying from 50 to 500-fold reduction). Although IFT88KD G0 cells display higher expression levels than Control G0 cells, they still show lower expression of these proliferative genes when compared to cycling cells (MB) (varying from 10 to 50-fold lower). Values represent mean ± s.e.m, N=3, * *p* < 0.01, ** *p* < 0.001, *** *p*< 0.0001

To identify networks among the genes that were differentially regulated in cilium knockdown cells at G0 we used the STRING (Search Tool for the Retrieval of Interacting Genes/Proteins) database (Szklarczyk et al., 2015). The up-regulated genes formed a tight network, corresponding to 2 major clusters representing cell cycle regulation and myogenic differentiation, and a minor cluster consisting of genes regulating ECM and related signaling (Fig. 3C). The down-regulated genes formed a more diffuse network of genes involved in the regulation of cell proliferation and response to stimulus (Fig. S5).

To gain further insight into the molecular pathways that may be altered in cells where ciliogenesis is blocked, we used GSEA analysis (Mootha et al., 2003; Subramanian et al., 2005), which showed an enrichment of genes related to cell cycle in the transcriptome of IFT88 knockdown cells (Fig. 3D). Notably, we observed an enrichment of genes that define a proliferation signature (pSig) when compared with earlier established gene sets defining quiescence and proliferation (Venezia et al., 2004) (Fig. 3D). Interestingly, the top ten gene sets with highest normalized enrichment score (NES) that were enriched in IFT88KD at quiescence conditions, represent genes that are involved in G2/M phases (Fig. 3E), which is consistent with the increased population of cells with 4N DNA content observed in the cell cycle profile (Fig. 2F). Accumulation of cells at G2/M phases might reflect non-ciliary roles of IFT88 (Delaval et al., 2011), or a requirement for IFT88 in G2/M entry, with blocked cells yet to exit the cell cycle at the time of harvest. Therefore, we analyzed a time course of entry into quiescence to reveal any shifts in cell cycle exit profiles. Surprisingly, cells lacking cilia displayed similar exit kinetics to control cells and showed no significant change in the proportion of either S phase cells (EdU^+^) or M phase cells (Histone3pS10^+^) (Fig. 3F) across the time course. Taken together, these results suggest that IFT88KD cells do not enter M phase of the cell cycle but are paused at G2.

To resolve the paradox of increased proliferative gene expression in cells displaying a non-proliferative phenotype, we analysed the expression levels of selected G2/M candidate genes that were up-regulated in knockdown cells. Control G0 cells showed 50-500 fold lower expression of transcripts encoding G2/M regulators AurA, Bub1, Cdc20, CenpF, FoxM1, Mcm3/5, and Plk4 compared to proliferating MB. Although there is increased expression of these G2/M marker genes in IFT88KD cells, the levels of these transcripts are still significantly lower than those observed in proliferating cells (10-50 fold lower) (Fig. 3G), suggesting that overall, the magnitude of altered expression may not achieve a threshold required for functional progression through G2 into M. Furthermore, GSEA analysis also revealed that the up-regulated genes in IFT88KD cells were involved in cell cycle checkpoints, in particular G2/M checkpoints involving DNA damage and mitotic spindle assembly, functions associated with the centrosome. Collectively, these results show that ablation of ciliogenesis leads to deviation from a canonical quiescence program, resulting in cells that are blocked at G2 rather than at G0, as defined by their transcriptional profile.

The second signature of up-regulated genes in IFT88KD cells represented myogenic regulators and muscle structural genes. Since the cilium is transiently induced during early differentiation and appears to be retained only in the minor population of non-differentiating reserve cells, the induction of myogenic genes in IFT88KD cells suggests that loss of the cilium de-repressed the differentiation program. The up-regulation of two gene networks representing opposing cell fates (cell cycle [G2/M] vs. differentiation) is consistent with the repression of both these states by signaling from the cilium.

### Myoblasts lacking cilia have reduced self-renewal potential

Induction of quiescence is associated with enhanced clonogenic self-renewal when cells are restored to proliferative conditions (Rumman et al., 2018). Cells with impaired ciliogenesis showed a deviation from canonical features of G0 such as expression of p27 and 2N DNA content. Therefore, we evaluated the self-renewal capability in IFT88KD myoblasts post-quiescence using colony formation (CFU assay) and found that knockdown myoblasts showed reduced CFU compared to control cells (Fig. 4A). The kinetics of cell cycle re-entry were also affected, with a significant decrease in EdU incorporation at 24 hours post re-plating (Fig. 4B). Interestingly, similar to the cell cycle effect, this reduction in self-renewal is also stage-specific, with no significant impairment in CFU formation in proliferating cultures (Fig. S6). These results show that even though IFT88 is expressed in both proliferating and quiescent cells, compromising its expression specifically in conditions where the cilium is normally elaborated leads to decreased self-renewal ability, consistent with the failure to enter a canonical quiescence program due to suppressed ciliary extension.

**Fig. 4:**
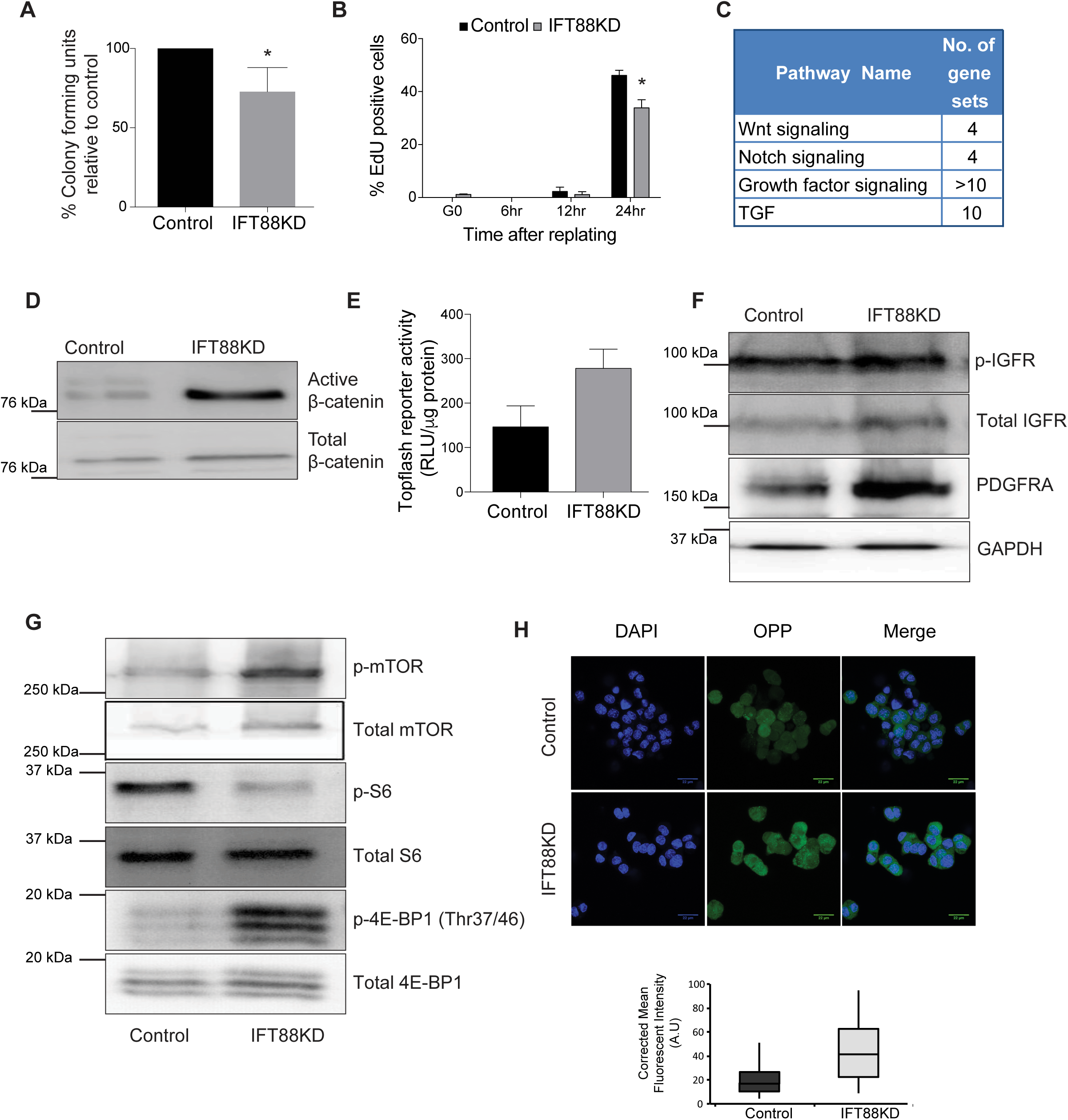
The altered quiescent state in IFT88KD is characterized by reduced self-renewal, deregulated reactivation kinetics and enhanced activity of proliferative signaling pathways. A. IFT88KD G0 myoblasts were harvested from suspension and replated at clonal density for analysis of colony forming potential (% CFU). IFT88KD myoblasts showed reduced self-renewal when compared to control cells Values represent mean ± s.e.m, N=4, \**p* = 0.0068. B. EdU incorporation was estimated during a time course of reactivation (6, 12, 24 hr) from quiescence (G0). Knockdown cells are able to return to the cell cycle, but at 24 hours when control cultures are restored to pre-suspension levels of DNA synthesis, IFT88KD cells showed significantly fewer cells in S phase. (Values represent mean ± s.e.m, N=3, *p-value<0.05). C-G Mitogenic signaling pathways are up-regulated by IFT88 knockdown. C. Gene sets related to pathways that are enriched in the IFT88KD G0 myoblast transcriptome. IFT88KD G0 transcriptome shows enrichment for genes related to cell signaling. D. Ablation of ciliogenesis results in increased expression of Wnt effector β-catenin. Immuno-blot (dephospho or active β-catenin) represents one of three independent experiments. E. Functional analysis: IFT88KD G0 cells show elevated signaling through the Wnt pathway. Quantification of increased Wnt signaling as indicated by higher TOPFlash activity (expressed as RLU/ μg protein). Values represent ± s.e.m, N=2, p-value=0.014. F. IFT88KD show elevated growth factor signaling in conditions of suspension arrest. Western blot analysis shows increased growth factor receptor protein expression for IGFR and PDGFRA, as well as increased activation of IGFR (p-IGFR). G. Activation of selective arms of translation regulatory pathways: IFT88KD cells at G0 conditions show elevated levels of phosphorylated (active) mTOR. However, there is divergent response in the key mTOR targets; while ribosomal protein S6 shows decreased phosphorylation, the translational repressor 4E-BP1 shows increased phosphorylation consistent with loss of repressive activity and indicating preferential activity through this effector node. H. IFT88KD myoblasts show increased global levels of protein synthesis. Protein synthesis was detected in Control and IFT88KD myoblasts which were placed under suspension arrest, using the Click-iT® Plus OPP Alexa Fluor® 488 Protein Synthesis Assay Kit (top panel), and the fluorescence levels were quantified in over 50 cells per sample (bottom panel).

**Fig. 5:**
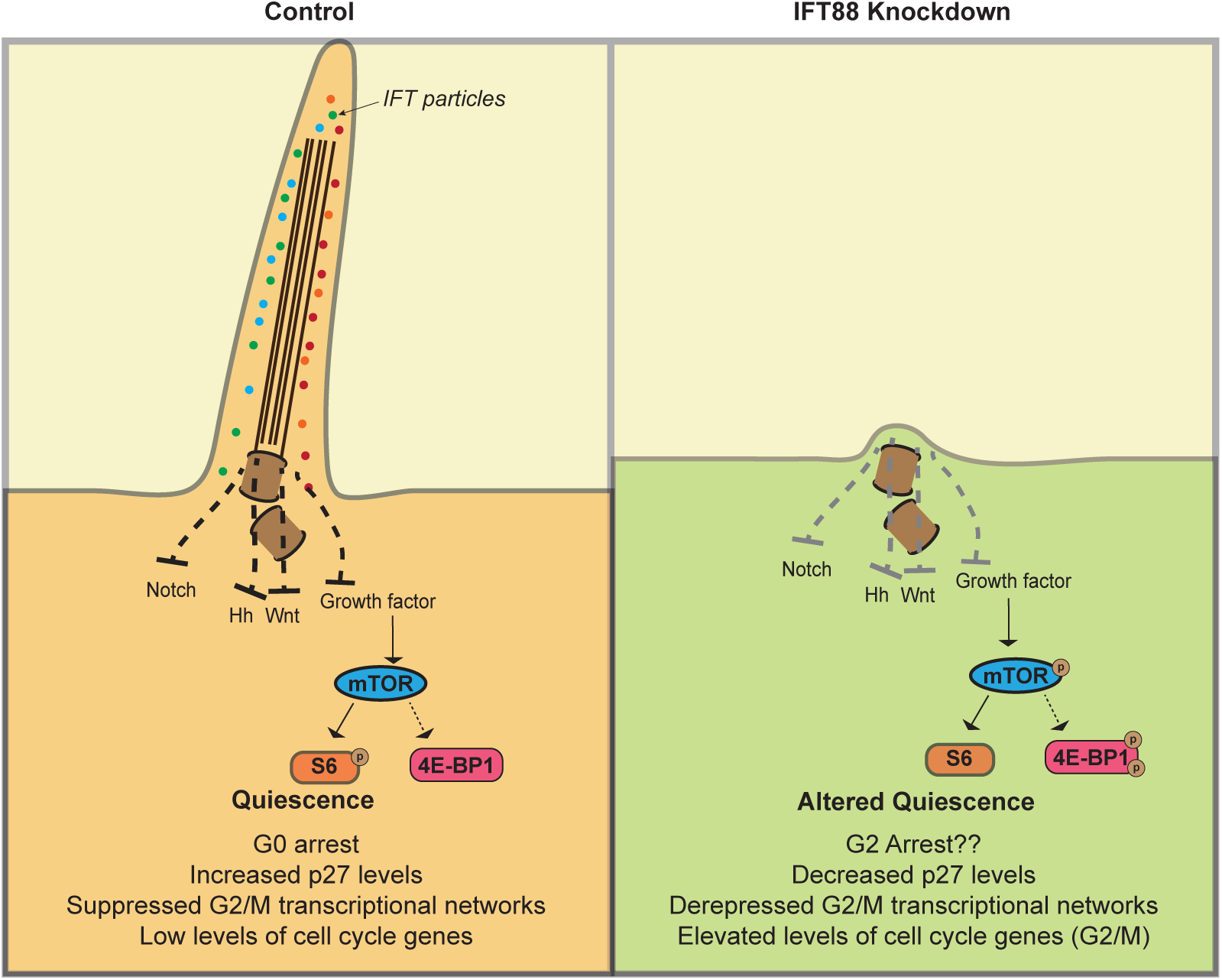
A working model for primary cilium function in myoblast quiescence. Control myoblasts entering suspension induced quiescence elaborate primary cilia, which work to suppress proliferative signaling in G0, whereas myoblasts where ciliogenesis is blocked enter an altered quiescent state characterized by arrest in G2 and activation of G2/M checkpoint genes.

### Myoblasts lacking cilia exhibit enhanced signaling activity

The cilium is a known sensory hub that harbors receptors for multiple signaling pathways (reviewed in (Basten and Giles, 2013; Singla and Reiter, 2006)). The enrichment of Wnt, Hh and mitogen receptors in the cilium is thought to enable growth factor-induced reactivation out of G0. In our culture model, quiescence is triggered by the abrogation of adhesion-dependent signaling pathways (Dhawan and Helfman, 2004; Milasincic et al., 1996). To elucidate the mechanism by which suppression of the primary cilium contributes to an altered quiescent program, we examined possible shifts in signaling cascades. Consistent with the notion of aberrant signaling, GSEA analysis of IFT knockdown cells showed an enrichment of genes related to cilium-dependent signaling pathways (Fig. 4C), including Notch, Hh, Wnt, and growth factor signaling. The primary cilium is known to show cell type- and condition-specific influences in either promoting or dampening the activity of these pathways (Wheway et al., 2018). Using a combination of reporter assays, qRT-PCR and western blot analysis, we assessed the activity of cilium-related pathways that have been previously implicated in G0 (Coller et al., 2006; Sachidanandan et al., 2002; Subramaniam et al., 2013). We detected enhanced signaling through 3 specific pathways in IFT88 knockdown myoblasts. Wnt signaling was elevated as evidenced by increased Wnt-TCF reporter activity (TOPflash), and induction of the transcriptional effector, active (dephospho) β-catenin (Fig. 4D, E). Increased levels of IGFR protein and phosphorylation were also seen, as well as increased levels of a key mediator of the G0-G1 transition, PDGFRA (Fig. 4F). These results suggest that under quiescence conditions, the primary cilium functions to dampen multiple growth factor signaling pathways.

We further examined whether the observed induction of upstream growth factor signaling events in knockdown cells led to enhanced activity at downstream signaling nodes. mTOR activity is an important integrator of growth factor signaling and functions by targeting protein synthesis (Fingar and Blenis, 2004). Suppression of ciliary extension and increased growth factor signaling resulted in increased mTOR phosphorylation (Fig. 4G), consistent with relay of an upstream signal such as the activation of PDGFR or IGFR. Two critical downstream targets of mTOR that directly affect translational activity are ribosomal protein S6 (rpS6), which is activated upon phosphorylation by S6 kinase (S6K), and 4E-BP1, a translational repressor which is inactivated upon phosphorylation (reviewed in (Showkat et al., 2014)). Interestingly, while rpS6 did not show an increase in phosphorylation (Fig 4G), IFT88KD myoblasts showed an increase in phosphorylation of 4E-BP1.

We next investigated whether the observed increase in the level of mTOR activity towards 4E-BP1 had phenotypic consequences i.e. increased translation. Indeed, we observed an appreciable increase in levels of protein synthesis in IFT88KD cells, as evidenced by the increased incorporation of OPP into newly synthesized proteins (Fig 4H). Taken together, this data suggests that the loss of the cilium channels mitogenic signals leading to translational control via one arm (4E-BP1) of the signaling pathway downstream of mTOR. 4E-BP1 phosphorylation has been shown to be specifically elevated during myogenic differentiation (Pollard et al., 2014), and during M phase in cycling cells (Velásquez et al., 2016), consistent with the enrichment of cells displaying these two expression profiles in IFT88 knockdown conditions. Thus, presence of the cilium in G0 maintains 4EBP1 in its translationally repressive unphosphorylated state and dampens overall protein synthesis.

In summary, this work shows the preferential extension of the cilium in muscle cells during reversible cell cycle arrest, but not during irreversible arrest/differentiation. Disruption of ciliation leads to an altered cell cycle distribution in IFT88KD cells, characterized by a block in G2 rather than G0, an altered transcriptome showing enhanced expression of regulators of G2/M checkpoints as well as enhanced expression of myogenic differentiation markers, reduced levels of classical quiescence indicators, aberrant re-entry and reduced self-renewal ability. Although suppression of ciliary extension correlated with increased activity of cilium-mediated signaling pathways, some downstream signaling nodes were not activated, likely accounting for the absence of enhanced proliferation. We conclude that cilium extension in quiescent myoblasts contributes to the acquisition of key features of G0 including the characteristic quiescence transcriptional program which features suppression of both cell cycle and myogenic genes, as well as dampened signaling to the translational machinery.

## Discussion

In this study, we have explored the function of primary cilia in skeletal muscle myoblasts and report a novel role for the organelle in the establishment of quiescence. Existing reports linking the cilium to the cell cycle have focussed on transition from G1 to S phase (Robert et al., 2007), as well as re-entry into the cell cycle from a quiescent state (Kim et al., 2011). These studies have shown that resorption and modification of ciliary length and function contribute to cell cycle progression. Similarly, signaling pathways such as Hh and PDGF that act through the cilium also contribute to regulation of cell cycle progression (reviewed in (Pan et al., 2013)) and asymmetric cell division (Jaafar Marican et al., 2016). Although the primary cilium is a quiescence-associated structure, there is surprisingly scanty evidence of the function of the cilium-centrosome axis in G0. Our study bridges this gap, and provides evidence for repressive signaling from the cilium in establishment and/or maintenance of quiescence.

We show that as in other cells, primary ciliogenesis in myoblasts is induced during cell cycle exit. While this concurs with the well-established reciprocal relationship between the cilium cycle and cell cycle, we find that ciliation is sustained in quiescence, but lost in terminal differentiation, suggesting a role in distinguishing these non-dividing states. Increased ciliogenesis is observed within 24 hours of receiving cell cycle exit cues, a time when expression of classical quiescence markers (such as p27) or differentiation markers (such as Myogenin) is still low. Thus, formation of the cilium occurs subsequent to receiving cell cycle exit cues (reduced adhesion or reduced mitogen availability), but prior to maturation of quiescent or differentiated states, implying a role for the signaling hub in this cell fate decision.

Our data also indicate that primary cilia mark only those cells that exit the cell cycle reversibly. Thus, in apparently homogeneous cultures induced to undertake myogenic differentiation, the primary cilium may help to select a sub-population (reserve cells) that resists differentiation cues and retains stem cell characteristics. We also observe heterogeneity in ciliation status within quiescent populations, suggesting that not all cells entering quiescence are equally capable of ciliogenesis. This may have implications for quiescent adult muscle stem cells, where some evidence exists indicating a correlation between heterogeneity and different propensities for activation (Collins et al., 2005). In conjunction with the finding that primary cilia are preferentially assembled on self-renewing MuSC in vivo (Jaafar Marican et al., 2016; this study), this observation provides interesting avenues for future exploration of ciliary roles in adult stem cell fate. The enhanced signaling activity in cilium-ablated cells may indicate that the cilium provides a zone of regulation for these pathways, possibly by physically localizing signaling intermediates or mechanical cues.

Strikingly, when cells with impaired ciliogenesis were placed under conditions of suspension arrest, while they did not attain quiescence, they also did not display continued cell proliferation as might be expected from increased mitogenic signaling. Instead, cells lacking cilia showed no increase in EdU incorporation (S phase), or phosphorylated H3 (M phase). This is also in contrast with an earlier report highlighting the role of IFT88 in mitotic spindle organization, which showed that IFT88 knockdown in HeLa cells resulted in mitotic catastrophe and delays (Delaval et al., 2011). The observed accumulation of cells in the G2 phase in IFT88KD suggest either an incomplete reversal/residual retention of ciliary function, or the presence of an unreported role for the cilium in G2/M checkpoint control. The dual role of the centrosome/basal body and the relationship of the centrosome-cilium cycle with the cell cycle suggests possible mechanisms. When IFT is impaired, while ciliary extension is compromised, the centrosome-basal body transition may not be affected. Thus, the centrosome is likely still docked as a basal body and may not be free for its mitotic spindle functions, resulting in G2/M accumulation of cells unable to traverse that block. Alternatively, the centrosome and basal body may have differential affinity for localization for cell cycle regulatory proteins and disturbing this balance may result in aberrant regulation of cell cycle progression, since cell cycle regulators such as Cyclin B1, AurA and Plk1 are reported to localize to the centrosome or basal body and this localization is essential for their function (Jackman et al., 2003; Spalluto et al., 2013). Since the centrosome alternates as a spindle organizer in M phase and the ciliary basal body in G0, it is possible that communication between the cilium and G2/M transcriptional networks is mediated by centrosomal components. Thus, the incomplete induction of G2/M transcriptional programs observed in the IFT88KD cells may reflect an additional requirement for transition to centrosomal function required for M phase entry.

The centrosome has also been linked with translational regulation of M phase regulators: Plk1was shown to co-localize with 4E-BP1 at centrosomes in mitotic cells, providing evidence for direct interaction and phosphorylation of 4E-BP1 by Plk1 (Shang et al., 2012). Interestingly, IFT88KD myoblasts induced to enter quiescence also show an increase in Plk1 expression as well as 4E-BP1 phosphorylation, accompanied by an increase in global protein synthesis. Considering emerging evidence of the functional association of translational machinery with the centrosome, our finding raises the possibility that the cilium-centrosome axis is involved in signaling between G2/M transcriptional programs and selective translation of 4E-BP1-dependent transcripts.

Our data shows that cells lacking cilia enter an alternate state of quiescence, characterized by a shift in transcriptional profile, as well as reduced ability to return to the cell cycle. These observations suggest that the primary cilium may also act as a transducer for quiescence signals and that the cilium-mediated establishment of the quiescence program is necessary for subsequent cell cycle re-entry and self-renewal. Disruption of primary ciliogenesis leads to a specific alteration of G2/M transcriptional networks in conditions of quiescence, suggesting a novel role for the control of mitotic progression by the cilium-centrosome axis. The observed halt at G2 in cilium-ablated cells indicates the requirement of signals in addition to the canonical G2/M transcriptome for transition through mitosis, such as mechanical cues, which may in turn also be sensed, transduced and regulated by the cilium/centrosome. While there is some evidence to support such non-ciliary roles for ciliary and centrosome proteins (Delaval et al., 2011; Jonassen et al., 2008; Kodani et al., 2013; Shang et al., 2012; Velásquez et al., 2016), our findings may also suggest that retention of the centrosome as a basal body is monitored, and that complete dismantling of the cilium/basal body complex is required for the G2/M transition.

Taken together, our results show that primary cilia are dynamically assembled when myoblasts receive cues for cell cycle exit, and strongly support the idea that the cilium is integral to the establishment of the quiescent state in skeletal muscle myoblasts, while also distinguishing reversible from irreversible arrest. The functions of the cilium in quiescence are mediated in part through the dampening of proliferative signaling, and suppression of cell cycle progression by influencing G2/M control mechanisms.

## Materials and Methods

### Cell culture

C2C12 myoblasts were obtained originally from H. Blau, Stanford University and a sub-clone A2 was derived in-house (Sachidanandan et al, 2002) and used for all experiments. Myoblasts were maintained in growth medium (GM; DMEM + 20% Fetal Bovine Serum (FBS) and antibiotics), ensuring that cells do not exceed 70-80% confluence before passaging. Differentiation was induced in low mitogen medium (DM: DMEM + 2% horse serum), for 5 days to form mature myotubes (MT). Synchronization in quiescence (G0) was induced by suspension culture of myoblasts for 48 hours in 1.3% methylcellulose medium prepared with DMEM containing 20% FBS, 10 mM HEPES, and antibiotics, as described (Sachidanandan et al., 2002). G0 cells were reactivated into the cell cycle by re-plating at a sub-confluent density in GM and harvested at defined times (30 minutes - 24hr) after activation.

### Immunofluorescence analysis and microscopy

#### C2C12 cells

Cells that were either plated on coverslips, or harvested from suspension cultures were simultaneously fixed and permeabilized in 2% PFA, 0.2% Triton X-100 in modified cytoskeletal buffer (CSK) (Gopinath et al., 2007) at 4**°**C for 15 min, to ensure detection of stable microtubules of the ciliary axoneme and depolymerization of dynamic cytoplasmic microtubules. Cells were incubated in blocking buffer (either 10% FCS, 0.2% Triton X-100 in CSK buffer or 5% BSA, 0.2% Triton X-100 in CSK buffer) for 1 hour to reduce non-specific labeling. Primary antibodies were diluted in blocking buffer. Specific antibody labeling was detected using fluorescence tagged secondary antibodies from Invitrogen. Details of primary antibodies used are as follows: Acetylated tubulin (SIGMA, T7451, 1:5000), Gamma tubulin (SIGMA, T6557, 1:1000), Acetylated tubulin (Abcam, ab125356, 1:1000), Ki67 (Abcam, ab1667, 1:200), Myogenin (Santa Cruz, sc-576, 1:100), Pax7 (AVIVA, ARP32742_P050, 1:1000), Pax7 (DSHB, 1:20), Pericentrin (Santa Cruz, sc-28147, 1:100). All washes were done in CSK buffer at RT. For suspension culture samples, immunostaining was performed as described earlier (Arora et al., 2017). Samples were mounted in aqueous mounting agents with DAPI. Images were obtained using either the Leica TCS.SP5-II AOBS, Leica TCS SP8 or Zeiss LSM 700 confocal microscopes. Minimum global changes in brightness or contrast were made, and composites were assembled using Fiji (ImageJ).

#### Single muscle fiber analysis

Animal work was conducted in the NCBS/inStem Animal Care and Resource Center. All procedures were approved by the inStem Institutional Animal Ethics Committees following norms specified by the Committee for the Purpose of Control and Supervision of Experiments on Animals, Govt. of India.

Single skeletal muscle fibers were isolated using a technique modified from (Shefer and Yablonka-Reuveni, 2005; Siegel et al., 2009). Briefly, the Extensor Digitorum Longus (EDL) muscle of 6-week old male C57 BL/7 mice was dissected out and treated with Collagenase Type 1 (Cat# LS4196 Worthington 400U/ml final concentration) for 1 hour at 37°C. The resultant dissociated muscle fibers were transferred into fresh DMEM and triturated to release individual muscle fibers, which were then washed through transfer to fresh medium, and then fixed with 4% paraformaldehyde for 15 min at RT. For immunostaining, single fibers were washed with PBS twice and mounted on charged slides (Cat#12-550-15, Fisher Scientific). The fibers were permeabilized with 0.5% Tween 20 in PBS for 1 hour at RT followed by blocking with 2 mg/ml BSA in PBS, for 1 hour at RT. Primary antibody incubations were performed overnight at 4°C. Secondary antibody incubations were for 1 hour at RT using fluorescently tagged antibodies from Invitrogen. All antibody dilutions were made in a solution of 1 mg/ml BSA in 0.25% Tween 20 in PBS. Antibody incubations were followed by three washes with blocking solution. 4’, 6-Diamidino-2-Phenylindole (DAPI) (Cat# 32670 Sigma) was used to stain the DNA. Images were acquired using Zeiss LSM 510 Meta confocal microscope.

### Western blot analysis

Lysates of adherent cultures or suspension cells were obtained after harvesting PBS washed cells by centrifugation, and resuspended in Laemmli sample buffer (2% SDS, 5% β 50 mM Tris-Cl pH 6.8) supplemented with Protease Inhibitor Cocktail and PhosStop (Roche) for isolation of total cellular protein. Protein amount in lysates was estimated using Amido Black staining and quantification of absorbance at 630 nm. Proteins were resolved on 8-12% Acrylamide gels, and transferred to PVDF membranes. The membrane was washed in 1X TBS with 0.1% Tween20 (TBST) and blocked in 5% blocking reagent (5% w/v nonfat milk in TBST) for 1 hour at room temperature, followed by incubation with primary antibody overnight at 4°C, then washed in 1X TBS + 0.1% Tween for 10 mins each followed by incubation with secondary antibody conjugated with HRP (Horse radish peroxidase) for 1 hour. After a brief wash with TBS-T for 10 minutes, ECL western blotting detection reagent for HRP was used for chemiluminescent detection using the ChemiCapt (Vilber Lourmat) gel documentation system. Details of antibodies used in this study are as follows: Cyclin A2 (Abcam, ab7956, 1:500), Cyclin B1 (Abcam, ab52187, 1:500), Cyclin D1 (Abcam, ab40754, 1:200), Cyclin E1 (Abcam, ab3927, 1:200), IFT88 (Proteintech, 13967-1-AP, 1:1000), p130 (Santa Cruz, SC-317, 1:200), p27 (BD, 610242, 1:2000), p21 (BD, 556430, 1:2000), MyoD (DAKO, M3512, 1:500), GAPDH (Abcam, ab9484, 1:2000), Myogenin (Santa Cruz, SC-12732, 1:1000), Active-Beta Catenin (Millipore, 05-665, 1:2000), Total-Beta Catenin (BD, 610153, 1:2000), PDGFR-alpha (Santa Cruz, SC-338, 1:500), Hes1 (Abcam, ab71559, 1:500), Phospho - mTOR (CST, 2971, 1:1000), Total - mTOR (CST, 2972, 1:2000), Phospho - rpS6 (CST, 4858, 1:1000), Total - rpS6 (CST, 2217, 1:2000), Phospho-4E-BP1 (Thr37/46) (CST, 2855, 1:1000), Total 4EBP1 (53H11) (CST, 9644, 1:1000), Phospho-IGF-I Receptor β (Tyr1135) (CST, 3918p, 1:1000), IGF-I Receptor β (CST, 9750p, 1:1000).

### Cell cycle analysis

Adherent cells were trypsinized, washed in PBS and pelleted by centrifugation. Suspension-arrested cells were recovered from methylcellulose by dilution with PBS followed by centrifugation as described earlier. Cell pellets were dispersed in 0.75 ml of PBS, and fixed by drop wise addition into 80% ice-cold ethanol with gentle stirring, following which they were briefly washed with PBS and resuspended in PBS with 40 μM of the DNA dye DR^TM^ (Cat. Μ No DR50050, Biostatus) per 10^6^cells. Cell cycle analysis was performed on a FACS Caliber® Cytometer (Becton Dickenson) using CelQuest® software and analyzed using FlowJo® software. At least 10,000 cells were acquired for each sample. Forward scatter and side scatter were used to gate cell populations and doublets were removed from analysis.

### Analysis of p27 levels

A stable C2C12 line expressing a p27 sensor (p27-mVenus, a fusion protein consisting of mVenus and a defective mutant of p27-CDKI, (p27K(-)) (Oki et al., 2014) was generated, transfected with either control or IFT88 siRNAs, and placed in suspension arrest. After recovery from methyl cellulose medium, these cells were assayed for proportion of mVenus positive cells by flow cytometry using a FACS Caliber cytometer (Becton Dickenson). CelQuest**®** software was used for acquisition and FlowJo® software was used to analyze the data.

### EdU incorporation Assay

Cells to be analyzed for DNA synthesis were pulsed with 10 μ EdU for 30 minutes, washed in PBS and fixed as described earlier. Samples were stored in PBS at 4°C till further processing. For staining, cells were permeabilized and blocked in PBS having 10% FBS and 0.5% TX-100 and labeling was detected using Click-iT® imaging kit (Invitrogen) as per manufacturer’s instructions. Samples were mounted in aqueous mounting agents with DAPI. Images were obtained using the Leica TCS SP8 confocal microscope. Minimum global changes in brightness or contrast were made, and composites were assembled using Fiji (ImageJ).

### OPP incorporation Assay

Cells to be analyzed for protein synthesis were pulsed with 20μ OPP for 30 minutes, washed in PBS and fixed as described earlier. Samples were stored in PBS at 4°C till further processing. For staining, cells were permeabilized and blocked in PBS having 10% FBS and 0.5% TX-100 and labeling was detected using Click-iT® imaging kit (Invitrogen) as per manufacturer’s instructions. Samples were mounted in aqueous mounting agents with DAPI. Images were obtained using the Leica TCS SP8 confocal microscope. Image intensity was calculated using Fiji (ImageJ) software, and corrected mean intensity (CMI = total intensity of signal – (area of signal × mean background signal)) was determined.

### Knockdown of gene expression using siRNA

C2C12 cells were cultured in growth medium until 80% confluent. The cells were then trypsinized and plated on to tissue culture dishes at appropriate cell density according to size of the dish. Approximately 16 hours post plating, the cells were transfected with siRNA) using Lipofectamine RNAiMAX (Invitrogen) as per manufacturer’s instructions. The cells were incubated with the RNA-Lipid complex for at least 18-24 hours following which they were used for further experimental analysis. Typically, siRNA-mediated knockdown was in the range of 50-90% at RNA level. Details of siRNA used in this study are as follows. For IFT88 knockdown: 5’-GCUGUGAACUCGGAUAGAU-3’ (Eurogentec) or siGENOME Mouse Ift88 (21821) siRNA-SMARTpool (Dharmacon, M-050417-00-0010); For Control: Scrambled siRNA (Eurogentec SR-NP001-001) or siGENOME Non-Targeting Pool #1 (Dharmacon, D-00126-13-20)

### Isolation of RNA from cultured cells and Quantitative real-time RT-PCR

RNA was isolated from cells using Trizol**®** (Invitrogen) according to manufacturer’s instructions, dissolved in nuclease-free water, quality checked by agarose gel electrophoresis and quantitated by spectrophotometry using the NanoDrop ND-1000UV-Vis spectrophotometer (NanoDrop Technologies, Wilmington, DE). cDNA was prepared from 1 μg total RNA using Superscript III (Invitrogen) and used in SYBR-Green (Applied Biosystems) based quantitative real time PCR analysis performed on an ABI 7900HT thermal cycler (Applied Biosystems) normalized to GAPDH levels. Fold change was calculated using normalized cycle threshold value differences2^−ΔΔct^. Primer sequences used in this study are as follows: GAPDH: forward 5’-AATGTGTCCGTCGTGGATCTGA −3’, reverse, 5’-GATGCCTGCTTCACCACCTTCT -3’; IFT88: forward, 5’-ATGTGGAGCTGGCCAACGACCT-3’, reverse, 5’-TGGTCGCAGCTGCACTCTTCACT-3’; FEZ1: forward, 5’-TGTACTTCGGTGCCAGGATG - 3′, reverse, 5’-GAGAGGGAAGGGTCCTCCAG-3’; PDGFRα: forward, 5’-GGTGGCCTGGACGAACAGAG-3’, reverse, 5’-GGAACCTGTCTCGATGGCACTC-3’; YPEL3: forward, 5’-TGCGGGCCAGCAGAAGAGCG-3’, reverse, 5’-GGAGCTAGGTCAGTCCCAGCCGT-3’; YPEL5: forward, 5’-GGCGCCACTGGTAGAGCATT-3’, reverse, 5’-CAGGATCACACGGCCTTCCT-3’; AURKA: forward, 5’-CGGTGCATGCTCCATCTTCC-3’, reverse, 5’-CTTCTCGTCATGCATCCGGC-3’; Cdc20: forward, 5’-CCGGCACATTCGCATTTGGA-3’, reverse, 5’-GTTCTGGGCAAAGCCGTGAC-3’; CenpF: forward, 5’-CAGCTGGTGGCAGCAGATCA-3’, reverse, 5’-GCTGGGAGTTCTTGGAAGGC-3’; Bub1: forward, 5’-TGCTCAGTAACAAGCCATGGAAC-3’, reverse, 5’-CCTTCAGGTTTCCAGACTCCTCC-3’; FoxM1: forward, 5’-ACTTTAAGCACATTGCCAAGCCA-3’, reverse, 5’-TGGCACTTGGGTGAATGGTCC-3’; Ki67: forward, 5’-TGGAAGAGCAGGTTAGCACTGT-3’, reverse, 5’-CAAACTTGGGCCTTGGCTGT-3’, Mcm3: forward, 5’-CCAGGACTCCCAGAAAGTGGA-3’, reverse, 5’-TGGAACACTTCTAAGAGGGCCG-3’; Mcm5: forward, 5’-TCAAGCGCCGTTTTGCCATT-3’, reverse, 5’-CTCACCCCTGCGTAGCATGA-3’; Plk4: forward,5’-GAAGGACTTGGCCACACAGC -3’, reverse, 5’-GAACCCACACAGCTCCGCTA-3’.

### Global transcription profiling using microarray

1 μg of total RNA isolated from Control or IFT88 KD cells was converted to cDNA using One-cycle labeling kit and amplified using IVT labeling kit following manufacturer’s instructions (Affymetrix). The normalized cRNA was fragmented, hybridized to mouse Affymetrix Genechips (430A 2.0), washed, stained and scanned as per Affymetrix protocols. The experiment was repeated with three different biological replicates and data analyzed using Affymetrix Gene Chip operating software (GCOS). All the CEL (cell intensity) files generated by Expression Console were then loaded into the R Bioconductor “affy” package for microarray analysis (Gautier et al., 2004). Briefly, the CEL files were normalized using Loess Normalization before proceeding to further analysis of differential expression between Control and IFT88KD samples for each specific cell state. Genes showing >1.5-fold differential expression with *P* ≤ 0.05 were selected and a subset validated by real time Q-RT-PCR. The raw data is available on the GEO database (Series GSE110742).

### Analysis of microarray data

#### STRING analysis

The lists of genes that were either up-regulated or down-regulated in IFT88KD myoblasts under G0 conditions were used as input to search for networks in the STRING database using default settings with only connected nodes being represented in the network diagram.

#### GSEA

Normalized expression values for Control (SCR) and IFT88KD were used as input dataset and the gene sets used were those from the C5 collection and those related to signaling, which are available on the Molecular Signatures Database (MSigDB), as well as a gene set identified from earlier published datasets (pSig) (Venezia et al., 2004). The enrichment analysis was carried out as described in the GSEA User guide (http://software.broadinstitute.org/gsea/doc/GSEAUserGuideFrame.html).

### Luciferase reporter assays for testing Wnt pathway activity

All assays were performed as described on stable clones expressing either the Super8X-TOP-flash (TCF site) construct (Veeman et al., 2003)(Veeman et al., 2003), TFC-1, or the FOP-flash (mutated TCF site) construct (Veeman et al., 2003), FFC-15. TFC-1 and FFC-15 clones were derived earlier (Aloysius et al., 2018; Subramaniam et al., 2013).

Luciferase activity was measured in lysates of TFC-1 and FFC-15 cells that were transfected with either IFT88 or negative control siRNAs. Assays were performed using the Luciferase Reporter Gene Assay Kit (Roche) to obtain data as relative light units (RLU). TCF activity was finally expressed as an Arbitrary Unit (AU), which was obtained after normalizing RLU to total protein estimated using BCA kit (Pierce).

### Statistical Analysis

Unless otherwise mentioned, all data represented are values derived from at least 3 biological replicates and is represented as mean ± s.e.m, analyzed using Student’s t-test, where p<0.05 was taken as significant.

## Acknowledgements

We thank Jay Gopalakrishnan (Cologne) and Krishanu Ray (TIFR), for helpful discussions, and Pavithra Chavali and Puja Singh for critical reading of the manuscript. We gratefully acknowledge Nandini Rangaraj (CCMB Advanced Imaging Facility), M.B. Madhavi (CCMB Microarray Facility), and CIFF at the Bangalore Life Sciences Cluster for imaging facilities.

## Author contributions

N.V. and J.D. designed experiments, interpreted data and wrote the manuscript; N.V., A.G. H.G., A.A., and N. Vyas performed experiments.

## Data availability

The raw data for the microarray analysis is available on the GEO database (Series GSE110742).

